# Gel-forming fibres differentially modulate inulin fermentation: A comparison of psyllium and methylcellulose in *in vitro* colonic models

**DOI:** 10.64898/2026.04.07.717018

**Authors:** Amisha A Modasia, Joshua E S J Reid, Alaa T Alhasani, Catherine Booth, Hannah Harris, Caroline Hoad, Penny A Gowland, Gleb E Yakubov, Maura Corsetti, Luca Marciani, Robin C Spiller, Frederick J Warren

## Abstract

Fermentable fibres such as inulin can support metabolic health but may exacerbate gastrointestinal symptoms in individuals with irritable bowel syndrome (IBS) due to rapid fermentation and gas production. The gel-forming fibre psyllium improves IBS symptoms, although the underlying mechanisms remain unclear. We hypothesised that fibre gelation alters fermentation by modulating microbial access to substrates. To test this, we compared psyllium with methylcellulose, a chemically modified, gel-forming fibre, to determine the effects of gelation on inulin fermentation.

Inulin alone or combined with psyllium or methylcellulose was fermented for 48 hrs in a colonic fermentation model inoculated with healthy human faeces. Gas production, metabolite profiles, microbial community composition and microbial localisation within fibre gels were assessed. Bioactivity of fermentation products was evaluated in STC-1 cells.

Psyllium co-fermentation significantly accelerated fermentation and enhanced production of metabolites, while methylcellulose had minimal effects. Psyllium maintained higher diversity and enriched polysaccharide-degrading taxa including *Bacteroides* and *Phoecaeicola* species, which were strongly associated with metabolic activity. Bacterial penetration into the psyllium matrix was observed but not into methylcellulose. Fermentation products from psyllium but not methylcellulose stimulated GLP-1 and 5-HT secretion in STC-1 cells.

These findings demonstrate that delayed-onset fermentable gel-forming fibres enhance microbial access to entrapped substrates, driving metabolic and hormonal responses.

## 2. Introduction

Dietary fibres (DFs) are structurally diverse carbohydrate polymers derived predominantly from plant-based foods. Although resistant to endogenous digestion by human enzymes (Stephen, Champ et al. 2017), DFs are differentially fermented by the gut microbiota within the colon, producing gas and metabolic products. These microbial metabolites, including short chain fatty acids (SCFAs), support host health by regulating epithelial integrity, modulating immune responses and influencing system metabolism (Blaak, Canfora et al. 2020, Cronin, Joyce et al. 2021, Gill, Rossi et al. 2021, Modasia, Spiller et al. 2026). The physiological outcomes of fibre consumption, therefore, depend not only on their chemical structure but also on how they shape microbial ecology and fermentation dynamics (Warren, Fukuma et al. 2018).

Despite the recognised health benefits of DFs, rapidly fermented DFs, particularly fermentable oligo-, di-, and mono-saccharides and polyols (FODMAPs), such as inulin (INU), can provoke unpleasant symptoms. Individuals with irritable bowel syndrome (IBS) often experience bloating, increased flatulence, and abdominal discomfort following FODMAP intake (Shepherd, Lomer et al. 2013, Gibson, Halmos et al. 2020). While fermentation of FODMAPs leads to luminal gas production in both healthy individuals and those with IBS, the total volume of gas produced is not necessarily greater in IBS. Rather symptoms are though to arise from visceral hypersensitive and altered central processing of gut-derived signals, resulting in heightened perception of luminal distention to which IBS patients are hypersensitive (Major, Pritchard et al. 2017). Consequently, normal physiological levels of gas and intestinal distention may be perceived as painful or uncomfortable in IBS. Although low-FODMAP diets can reduce symptoms, they are difficult to adhere to and may negatively impact microbial diversity and long-term gut health (Staudacher, Lomer et al. 2012). Strategies that slow fermentation and shift it towards the distal colon may alleviate these symptoms while preserving the health benefits of DF.

One promising approach involves combining rapidly fermentable fibres with viscous, gel-forming fibres. Psyllium (PSY) is a highly branched, non-starch polysaccharide that forms a viscoelastic gel upon hydration. It has been shown to slow INU fermentation *in vivo*, attenuating gas production (Major, Murray et al. 2018, Gunn, Abbas et al. 2022) whilst preserving SCFA production (Alhasani et al., 2024). Furthermore, we recently compared PSY to a preparation of methylcellulose (MC) a gel-forming cellulose derivative which resists bacterial degradation (Campbell and Fahey 1997, So, Yao et al. 2021) in delaying in delaying colonic fermentation of INU (Reid, Alhasani et al. 2026). While MC produced a smaller and non-significant reduction in initial breath hydrogen production compared to PSY, we speculated that this was due to MC not delaying orocaecal transit as observed for PSY. Slower fermenting fibres such as PSY confer health benefits beyond SCFA production by modulating gut motility, barrier integrity, microbiota composition, and inflammation, highlighting their therapeutic potential for symptom management without gas-related discomfort (Modasia, Spiller et al. 2026).

Fermentation dynamics are also closely linked to host gut–microbe communication. The SCFAs produced during fermentation stimulate enteroendocrine cells (EECs), specialised chemosensory cells that detect luminal nutrients and microbial metabolites. They respond by releasing peptide hormones such as glucagon-like peptide-1 (GLP-1) and serotonin (5-HT) that play key roles in regulating functions including intestinal secretion and motility (Gribble and Reimann 2016). Altered EEC density and signalling have been reported in patients with IBS (Dunlop, Coleman et al. 2005, Dunlop, Hebden et al. 2006), while DFs and the gut microbiota have been shown to influence EEC populations (Mazzawi and El-Salhy 2016, Mazzawi and El-Salhy 2017, Modasia, Parker et al. 2020). However, it remains unclear whether changes in the kinetics of fermentation, particularly when altered by fibre gelation, translate to microbial metabolic alterations and downstream gut hormone responses.

To address this gap, the present study investigates how gel-forming fibres modulate microbial fermentation of INU and the downstream effects on gut–microbe–host signalling. We compare PSY (a fermentable gel-forming fibre) with MC (a non-fermentable, gel-forming fibre). By examining a fermentable versus a non-fermentable gel matrix, we aimed to disentangle the role of gelation from fermentability in modulating microbial fermentation kinetics.

Using an *in vitro* anaerobic batch fermentation model inoculated with healthy human stool, we assessed gas kinetics, SCFA production, and microbial compositional changes over 48 hours. In parallel, we evaluated whether fermentation supernatants modulate hormone expression and secretion in the intestinal secretin tumour cell line (STC-1) (Fig. 1). Together, these analyses provide mechanistic insight into how physical properties of DFs influence microbial fermentation and their subsequent effects on gut signalling in the host. These findings may contribute to the development of more tolerable strategies for incorporating fermentable fibres into the diets of individuals with IBS.

**Figure 1.**
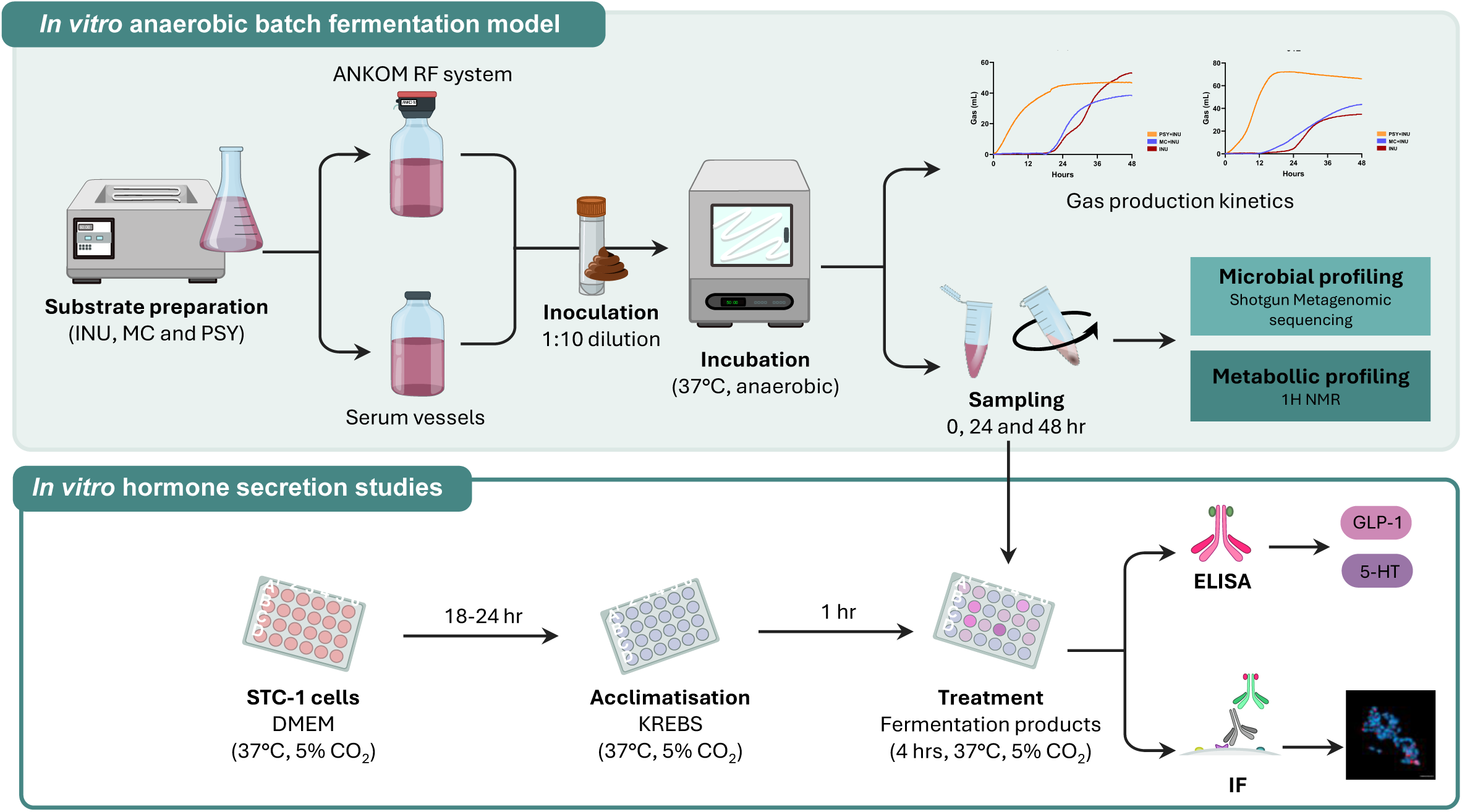
Schematic overview of the *in vitro* experimental design. Anaerobic batch fermentation was conducted using stool samples from healthy human donors (n=20) to assess colonic fermentation of inulin with or without the addition of gelling fibres, psyllium and methylcellulose. Gas production kinetics were measured using the ANKOM RF system. Samples were collected at 0, 24, and 48 hours for microbial and metabolic profiling. To evaluate the impact of fibre fermentation products on hormone responses, STC-1 cells were treated with fermentation supernatants. Hormone expression and secretion were analysed by immunofluorescence and ELISA, respectively. *Abbreviations: INU (inulin); MC (methylcellulose); PSY (psyllium); ELISA (enzyme-linked immunosorbent assay); 5-HT (serotonin); GLP-1 (glucagon-like peptide 1); ELISA (enzyme-linked immunosorbent assay); IF (immunofluorescence).* Graphics created by A Modasia (Inkscape 2.0).

## 3. Methods

### 3.1. Study population

Healthy volunteers (n=30) were recruited to a single centre, single-blinded, randomised, three-way crossover dietary intervention study conducted from April to August 2023 at the Nottingham Clinical Research Facility (N-CRF), Queens Medical Centre, Nottingham, UK (Reid, Alhasani et al. 2026). The study design was reviewed and approved by the Research Ethics Committee of the Faculty of Medical and Health Sciences in July 2022 (FMHS 19-0622) and registered at www.clinicaltrials.gov (ID: NCT05911347).

For FISH experiments, faecal samples were provided as part of an ongoing human study, “QIB Colon Model Study”. The study was approved by the local Quadram Institute Bioscience Human Research Governance committee (IFR01/2015) and by the London-Westminster Research Ethics Committee (15/LO/2169). The trial was registered at clinicaltrials.gov (NCT02653001). All participants provided signed informed consent prior to donating samples. The trial was registered at www.clinicaltrials.gov (ID: NCT02653001).

### 3.2. Stool sample collection

Participants (n=30) were provided with a stool collection kit and asked to collect double bagged samples with an added Oxoid AnaeroGen 2.5L Sachet (Thermo Scientific) to reduce oxygen atmosphere and store them in their domestic freezer until they brought them to the researcher on their first study day visit. Once received on site, samples were transferred to - 80°C at the Biomedical Research Unit prior to being sent to the Quadram Institute at Norwich Research Park for further analysis of their fermentation potential.

### 3.3. *In vitro* batch fermentation

Gas production of the test fibres (INU, co-fermentation of INU with PSY (INU+PSY) and MC(INU+MC)) was measured using single-stage anaerobic fermentation models (Williams, Bosch et al. 2005) with the ANKOM RF gas production system (ANKOM Technology, Macedon, New York, USA). Gas production from the test substrates was calculated using previous methods (Rotbart, Yao et al. 2017, Alhasani, Modasia et al. 2024). The data are reported as the cumulative gas volume (mL) produced during fermentation from 0-48 hr.

#### 3.3.1. Media preparation

Basal media was prepared as described by Williams et al., (Williams, Bosch et al. 2005). In brief, per 125 mL vessel, 38 mL basal solution with 3.5 mL of phosphate vitamin and buffer solution, and 0.5 mL of reducing solution were mixed and pH was adjusted to 6.8-7. Vessels were kept anaerobic under a constant stream of CO_2_.

#### 3.3.2. Substrate preparation

Vessels containing media were pre-warmed to 80°C, and substrates were added to the media with continual magnetic stirring. Per vessel, 0.5 g INU (Orafti HP), 0.5 g INU with 0.5 g PSY (INU+PSY) or 0.5 g INU with 0.5 g MC (INU+MC; 1:1 w/w mixture of Benecel A4M and Benecel MX) were added. Fermentation bottles containing no substrate were included as a control to remove any discrepancies in gas produced from sources other than the test substrates. For vessels containing MC, an additional step of cooling to 37°C using an ice bath was carried out after heating to 80°C and the addition of substrates. Vessels were then sealed, purged with CO_2_ and left overnight at 37°C with gentle agitation (∼80 rpm) prior to inoculation.

#### 3.3.3. Faecal slurry preparation and inoculation

Faecal samples were frozen at -80°C prior to use. Each faecal sample was defrosted at 4°C overnight, diluted in pre-reduced sterile PBS (10 % wt/v) and homogenised by vortexing with glass beads. Each substrate was fermented in duplicate per volunteer faecal sample. Fermentation vessels were seeded with faecal samples from individuals from the human study. For gas experiments, 12 participants were selected based on the availability of sufficient faecal material, as the colon fermentation model (ANKOM gas method) requires larger volumes of media and consequently greater quantities of faecal inoculum than standard *in vitro* protocols. Following inoculation with 1.5 mL faecal slurry, vessels were purged with CO_2_ and incubated at 37°C. Gas production was automatically measured every 5 min using the ANKOM RF system. Data are reported as cumulative gas volumes (mL) produced during fermentation, averaged from 12 individuals and measured in duplicate per individual/substrate.

#### 3.3.4. Sampling

For sampling, batch fermentation models were set up in parallel as described above in 125 mL serum bottles and 1 mL of sample was extracted using a 18G needle and syringe at 0, 24 and 48 hr. Samples were centrifuged at 10, 000 g for 5 min. Pellets and supernatant were stored at -80°C.

### 3.4. Metabolomics

NMR buffer was prepared using D_2_O as the solvent and the following concentrations of reagents: 21.7 mM NaH_2_PO_4_, 82.7 mM K_2_HPO_4_, 8.6 mM NaN_3_, 1.0 mM 3-(trimethylsilyl)-propionate-d4 (TMSP). Acetate, propionate, butyrate, succinate, and lactate concentrations were quantified in the contents of *in vitro* batch fermentation vessels following 24 hr fermentation using ^1^H NMR spectroscopy. Samples were thawed and prepared by mixing 400 µL of sample with 200 µL of NMR buffer and pipetted to NMR tubes (Norell® Standard Series™, 5 mm). The ^1^H NMR spectrum was acquired for 64 scans, a spectral width of 12, 500 Hz and an acquisition time of 2.62 s. Acetate, propionate, butyrate, succinate, and lactate concentrations were quantified using the Chenomx NMR Suite v10.0.

### 3.5. DNA extraction and metagenomics

Microbial pellets collected from *in vitro* fermentation were thawed at room temperature (21°C) and DNA extracted using the Maxwell® RSC Faecal Microbiome DNA Kit (Promega) according to recommendations of the manufacturer.

Library preparation was carried out using a modified version of the Illumina Nextera DNA Flex Library Prep Kit, as described in (Alhasani, Modasia et al. 2024). Metagenomic sequencing was carried out using three 10B lanes of an Illumina NovaSeq X Plus (PE15025B) instrument by Novogene (Cambridge, UK). This yielded an average of 10GB of data per sample.

### 3.6. Bioinformatic analysis

The resulting reads were analysed using BioBakery 3 Tools that include Kneaddata for processing and MetaPhlAn4 for taxonomic analysis (Blanco-Míguez, Beghini et al. 2023). Data visualisation was carried out using the Phyloseq (McMurdie and Holmes 2013) and Phylosmith R packages (Smith 2019).

Alpha and beta diversity were assessed using Shannon and Simpson indices across substrates and timepoints. Community composition was explored using distance based-redundancy analysis (dbRDA) constrained by participant. Associations between Shannon diversity and total gas production at 24 hr were assessed using Spearman correlation. Relative abundance plots were generated for taxa significantly associated with substrate and metabolic outputs based on mixed-effects modelling.

### 3.7. Fluorescent *in situ* hybridisation

*In vitro* fermentation of PSY and MC alone without INU (4% w/v) was conducted using methods described in 3.4 using stool samples from 3 healthy donors (QIB Colon Model Facility. The Fluorescent *in situ* hybridisation (FISH) staining method was adapted from Gorham et al., 2016 and Harris et al., 2023 (Gorham, Williams et al. 2016, Harris, Pereira et al. 2023). PSY and MC gel samples were collected at 24 hr fermentation, mounted into OCT embedding matrix and frozen using a dry ice-ethanol bath. Samples were stored in dry ice or at -80°C prior to sectioning on the cryotome. Samples were cryosectioned at -40°C and 10 μm slices were taken and mounted on SuperFrost™ microscope slide.

A circle was drawn around the sample using a PAP pen, and samples fixed in 4% paraformaldehyde for 30 min at 21°C. Slides were washed in PBS, then dehydrated by immersing for 3 min sequentially in 50%, 80% and 100% ethanol and air dried. The probes were then added at 50 ng/μL in hybridisation buffer solution (per 20 ml 3600 μL 5M NaCl, 400 μL 1M Tris-HCl (pH 8.0), 600 μL formamide, 9980 μL double-distilled water (ddH2O) and 20 μL 10% SDS) was added and incubated in dark for 1 hr at 50°C with humidity. The probes used were Eub338-Cy5 probe (probe sequence 5’-GCT GCC TCC CGT AGG AGT -3’; (Amann, Binder et al. 1990) and Bac303-TxRed (probe sequence 5’-CCA ATG TGG GGG ACC TT -3’; (Manz, Amann et al. 1996)). The samples were then washed with pre-warmed wash buffer solution twice for 15 min (per ml 12.8 μL 5M NaCl, 20 μL 1M Tris-HCl [pH 8.0], 10 μL 0.5M EDTA [pH 8.0], 96.2 μL ddH2O and 1 μL 10% SDS). Slides were washed in dH2O, air dried and counterstained for 30 min at 21°C with 1:100 Calcofluor White (Sigma) and mounted using FluorMountG and sealed with coverslip and nail varnish.

### 3.8. STC-1 cell culture experiments

The intestinal secretin tumour cell line (STC-1 cells) was purchased from ATCC (CRL-3254). Cells were cultured and maintained in Basal medium containing Dulbecco’s Modified Eagle Medium (DMEM; ATCC) supplemented with 10% FBS (ATCC) and 1% penicillin/streptomycin at 10,000 U/mL (ThermoFisher). Cells were incubated in a humidified CO_2_ incubator at 37 °C. All the experiments were performed at 80% confluency and passage number between 18-20. For hormonal secretion, cells were seeded in 12-well chamber slides (THISTLE SCIENTIFIC) at a density of 1.5 x10^4^ cells/well and incubated for 18-24 hrs prior to the experiment. Cells were acclimatised in KREBS buffer for 1 hr at 37°C in 5% CO_2_. Following this, cells were incubated with 0.22µm filter sterilised test substrates diluted 1:10 in KREBS buffer (Thermo Fisher) for 4 hrs. No substrate and a SCFA mixture (containing 10 mM acetate, propionate and butyrate at a ratio of 60:20:20, respectively) were used as negative and positive controls, respectively. Halt Protease and Phosphatase Inhibitor (Thermo Fisher) was added to each well to protect against gut hormone degradation and supernatants were aspirated, centrifuged (900 g x 5 min) and aliquoted at -80°C prior to hormone analysis.

#### 3.8.1. Immunofluorescence

Immunocytochemistry was performed to evaluate the changes in GLP-1 and 5-HT expression in STC-1 cells. Following treatment, cells were washed with PBS and fixed with PBS/4% paraformaldehyde for 20 min at room temperature. After this, cells were washed with PBS and membranes permeabilised with 0.25% Triton-X100 in PBS (MERCK LIFE SCIENCES UK LIMITED) for 15 min and incubated with either 10% goat or donkey serum (MERCK LIFE SCIENCES UK LIMITED) for 30 min. Cells were then treated with either Anti-GLP-1 (Bioss Antibodies, bs-0038R) and anti-serotonin (Abcam, AB66047) for 1 hr at room temperature and washed again with PBS, incubated with secondary antibodies (goat anti-rabbit or donkey anti-goat; Invitrogen Alexa Fluor™ 594) for 30 mins at room temperature, followed by Hoechst 33342 (Abcam) for nucleus staining. Finally, cells were washed in PBS and mounted using Invitrogen™ ProLong™ Diamond Antifade mountant (ThermoFisher). Cells were imaged using a Zeiss Axio Imager M2 microscope equipped with 40x/air objective and Zen blue software (Zeiss). All images were captured at 40X magnification.

Fluorescence was recorded at 405 nm (blue) and 594 nm (red) and intensity quantified using the sum fluorescent pixel intensity of the field of view (FOV) normalised to nuclear counts using a macro written in Image J/FIJI v1.52p. A minimum of 10 FOV images were used for any subsequent quantification.

#### 3.8.2. Gut hormone secretion

Gut hormone release (GLP-1 and 5-HT) from the STC-1 cells was quantified using ELISA. Supernatant from STC-1 cells collected following treatment was used to perform ELISA with commercial kits following manufactures’ protocols (EZGLP1T; MERCK LIFE SCIENCES UK LIMITED and STJE0006586; St John’s Laboratory).

### 3.9. Statistical analysis

All data in box-whisker plots are presented as mean ± standard error of the mean (SEM) with the indicated sample sizes. Box plots represent the first quartile, median and third quartile, with whiskers representing the minimum and maximum. Statistical analyses were preformed using GraphPad Prism (v10) and RStudio (2023.06.1 Build 524). Unless otherwise stated, a value of *p* < 0.05 was considered statistically significant. Linear mixed-effects models were implemented in RStudio using the lme4 package to analyse SCFA data, accounting for repeated measures where appropriate. For comparisons involving multiple groups, one-way or two-way ANOVA followed by Tukey’s multiple comparisons test was performed using GraphPad Prism, as indicated in the relevant figure captions.

## 4. Results

### 4.1. Psyllium increases the rate of inulin fermentation *in vitro*

Co-fermentation of INU with PSY (INU+PSY) significantly increased the rate of gas production compared to INU fermented alone, indicating that PSY modifies fermentation kinetics. A rapid increase in gas accumulation occurred within the first few hours of fermentation (Fig. 2A). Quantitative analysis confirmed a significant (p<0.0001) increase in total gas peak area at both 0-24 hr (Fig. 2B; 609.5±11.4 mL vs 78.0±10.0 mL of INU+PSY and INU, respectively) and 0-48 hr (Fig. 2C; 3152.0±35.1 mL vs 1914.0±35.0 mL of INU+PSY and INU, respectively), indicating a sustained increase in fermentation activity over time.

**Figure 2.**
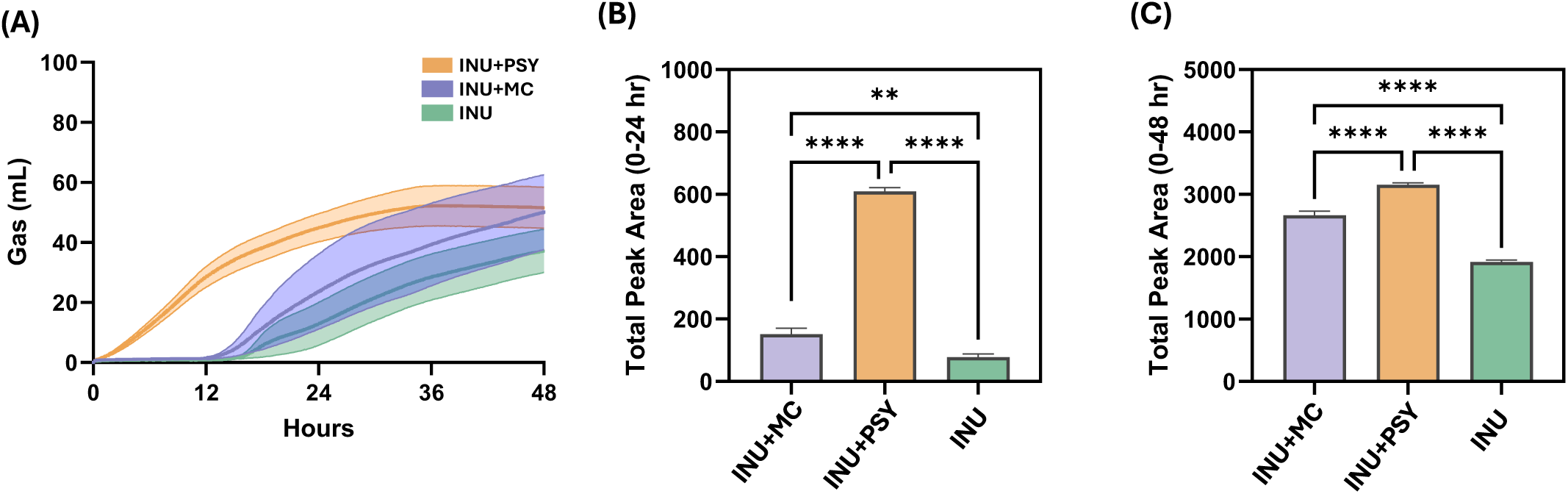
Psyllium increases rate of inulin fermentation *in vitro.* (A) Gas production kinetics assessed using ANKOM RF system during anaerobic colonic fermentation of inulin (INU), inulin + psyllium (INU+PSY) and inulin + methylcellulose (INU+MC) using stool samples from healthy volunteers (n=12) over a 48 hr period. Total peak area calculated for (B) 0-24 hr and (C) 0-48 hr fermentation. Data are represented as mean ± SEM Statistical significance calculated using ordinary one-way ANOVA with Tukey’s multiple comparisons test (GraphPad Prism v10). Values of *p<0.05, **p<0.01, ***p<0.001 and ****p<0.0001 were considered statistically significant.

Although MC co-fermentation (INU+MC) also modulated INU fermentation kinetics, its effects were less pronounced than with INU+PSY. INU+MC resulted in a modest significant increase in gas production compared to INU alone; however, total gas production peak area at 0-24 hr (Fig. 2B; 151.5±19.0 mL) and 0-48 hr (Fig.2C; 2663.0±68.4 mL) were significantly (p<0.0001) lower than those observed with PSY.

The accelerated fermentation observed during INU+PSY fermentation was not attributed to PSY fermentation itself. Gas production associated with PSY remained minimal during the early phase of incubation, with appreciable increases occurring only after ∼20 hrs, consistent with delayed kinetics. In contrast, MC remained non-fermentable, with no notable increases in gas production throughout incubation (Fig. S1).

Together, these findings indicate that PSY not only accelerates the onset of INU fermentation *in vitro* but also enhances the overall extent, whereas MC exerts a weaker modulatory effect on fermentation kinetics. We also observed that while PSY fermentation produces a foam with many small bubbles this was not seen with MC even at 48 hours.

### 4.2. Enhanced SCFA production is observed following co-fermentation with psyllium and inulin

Co-fermentation of INU with PSY (INU+PSY) resulted in increases in SCFA production (Fig. 3). Notably, concentrations of propionate and butyrate were significantly (p<0.01) elevated compared to INU alone. The most pronounced increase was observed for butyrate (Fig. 3C; 1.5±0.3 mM vs 0.3±0.1 mM for INU+PSY and INU, respectively), with propionate also showing a moderate but significant rise (Fig. 3B; 1.3±0.3 mM vs 0.4±0.1 mM for INU+PSY and INU, respectively). Increased acetate production was observed but this did not reach statistical significance (Fig. 3A; 11.7±1.6 mM vs 5.0±0.8 mM for INU+PSY and INU, respectively; p=0.06).

**Figure 3.**
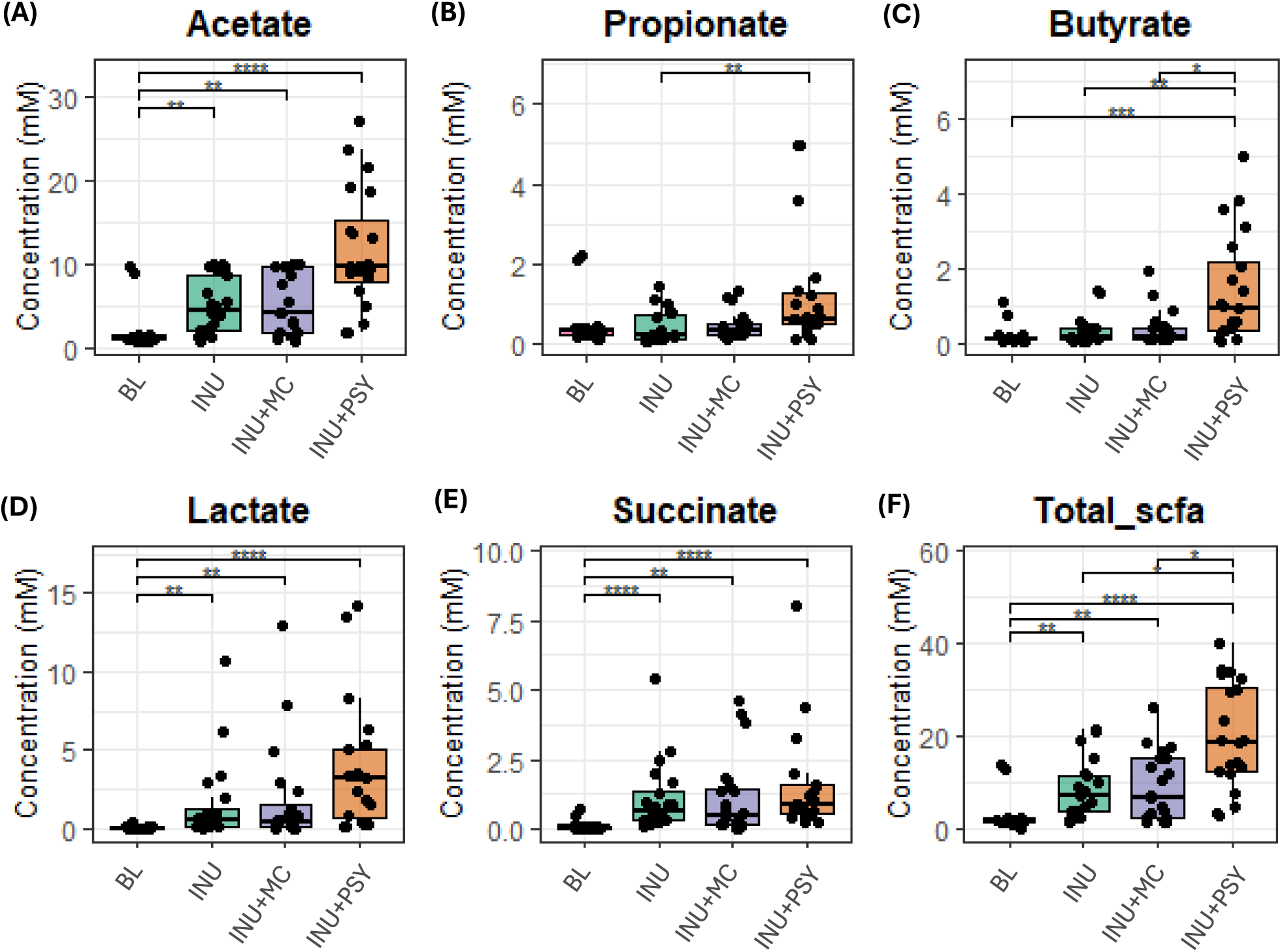
Psyllium enhances SCFA production during *in vitro* fermentation. (A) Acetate, (B) propionate, (C) butyrate, (D) succinate, (E) lactate and (F) total SCFA (acetate, propionate and butyrate) concentrations (mM) in media analysed following 24 hr fermentation of inulin (INU), inulin + psyllium (INU+PSY), inulin + methylcellulose (INU+MC) and blank (no substrate control) using 1H NMR (n=20). Statistical significance calculated using linear mixed modelling (RStudio). Values of *p<0.05, **p<0.01, ***p<0.001 and ****p<0.0001 were considered statistically significant.

While cumulative gas production was substantially higher during INU+PSY fermentation, only a weak association was observed between gas area under the curve (AUC) and total SCFA production at 24 hr (R² = 0.03; Fig. S2B), indicating that gas output was not a strong predictor of SCFA yield. Notably, the order-of-magnitude increase in gas production corresponded to only an approximate two-fold increase in total SCFA concentrations, suggesting that PSY alters fermentation efficiency and/or metabolic activity rather than simply increasing overall SCFA output in proportion to gas production.

In addition to the major SCFAs, levels of the intermediary metabolite lactate were also elevated across all substrates. While lactate elevation showed a strong trend levels were 2.5-fold higher with INU+PSY (Fig. 3D; 3.8±0.9 mM), but this difference did not reach statistical significance compared to the other substrates.

### 4.3. Psyllium shapes microbial diversity and community structure during co-fermentation with inulin

Across all substrates, microbial diversity declined by 24 hr of fermentation, a common feature of *in vitro* colonic fermentations, where early fermentation phases driven by the rapid expansion of fast-growing taxa result in reductions in community evenness (Fig. 4A). INU and INU+MC show the largest decreases (p<0.01) in Shannon diversity compared to blank. In contrast, INU+PSY retained significantly higher Shannon diversity at 24 hr (p<0.01) compared to INU and INU+MC and 48 hr (p<0.01) when compared to INU suggesting a more even or less selective microbial response and balanced microbial community as fermentation progresses. A similar pattern is observed for Simpson diversity (Fig. 4B), indicating less dominance and more community balance with INU+PSY co-fermentation.

**Figure 4.**
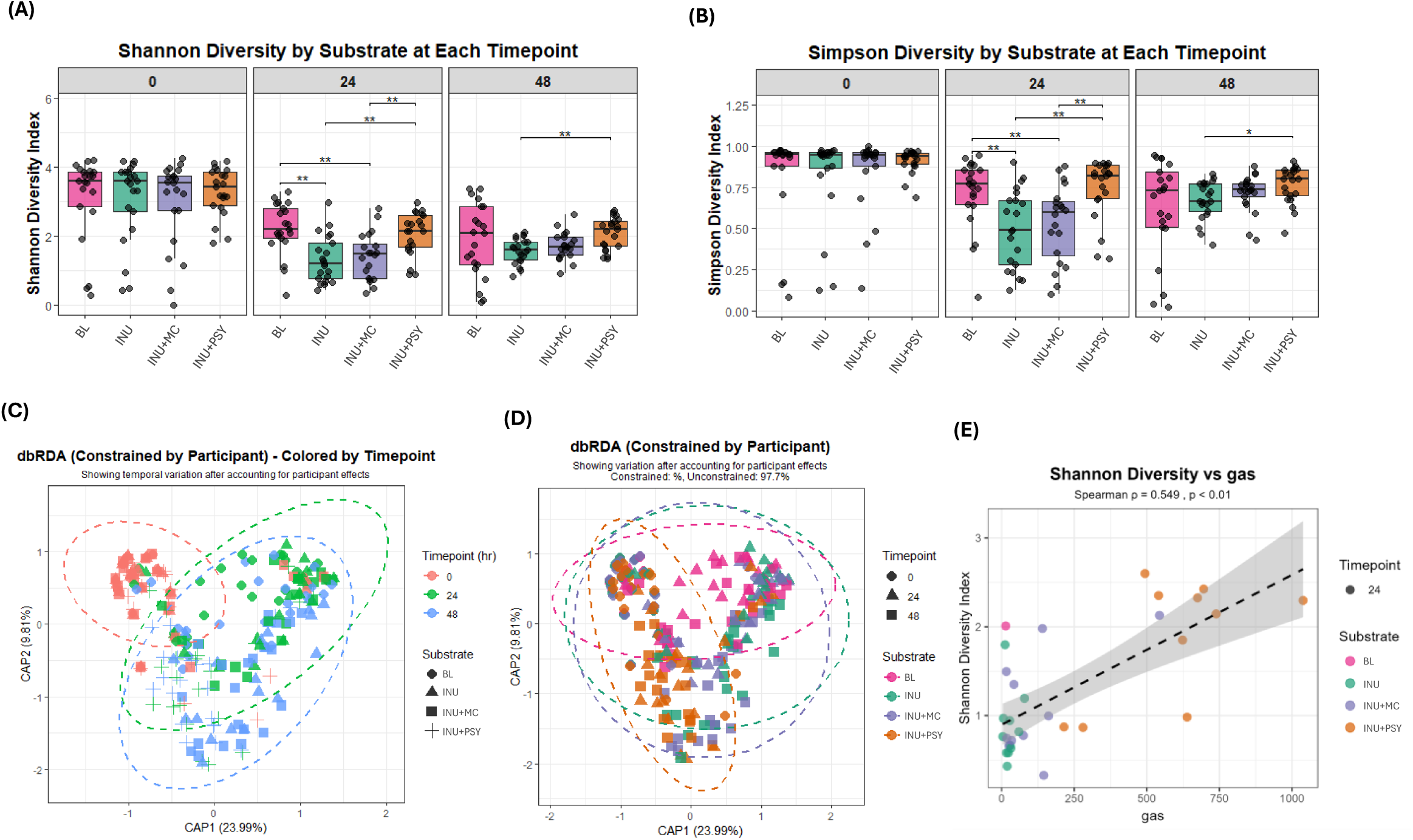
Drivers of microbiota diversity and community structure during *in vitro* fermentation. (A) Shannon diversity and (B) Simpson diversity by substrate (Inulin (INU); inulin + psyllium (INU+PSY); inulin + methylcellulose (INU+MC) and no substrate (BL)) and timepoint (0, 24 and 48 hr). Statistical differences in overall diversity were tested using Kruskal-Wallis rank sum test (*p<0.05 and **p<0.01). dbRDA plot constrained by participant coloured by (C) timepoint and (D) substrate. (E) Association between Shannon diversity and total gas production at 24 hr fermentation (Spearman ρ=0.549, p <0.01).

Community structural differences were further visualised using distance-based redundancy analysis (dbRDA) constrained by participant to show variation after accounting for inter-individual effects. After accounting for inter-individual variability, microbial community composition shifted systematically over time (Fig. 4C), with fibre substrate further modulating the direction and extent of this response (Fig. 4D). PERMANOVA pairwise analyses (Figure S3) indicated no significant differences between treatments at 0 hr. At 24 hr, however, the INU+PSY fermentation differed significantly from all other treatments (BL, INU, and INU+MC; p < 0.01), indicating an early and distinct shift in community composition driven by PSY, while INU and INU+MC remained similar to baseline and to each other. By 48 hr, additional differences were observed, with both INU (p < 0.05) and MC (p < 0.01) also differing significantly from BL. Overall, fermentation-driven divergence increased over time, with INU+PSY exerting the strongest and most persistent effect, whereas INU and INU+MC induced more gradual compositional shifts.

A positive linear association was observed between total gas production at 24 hr and microbial Shannon diversity (Spearman ρ = 0.549, p < 0.01; Fig. 4E). Higher gas output corresponded to increased microbial diversity, suggesting that more metabolically active fermentations supported a broader distribution of taxa. While diversity-gas relationships varied between substrates, the overall trend indicates that elevated fermentative activity is linked to increased community richness and evenness.

### 4.4. Psyllium selectively enriches fibre-degrading taxa and links microbial profiles to SCFA production

Substrate-specific changes in microbial taxa are shown in Figure 5. INU+PSY fermentation induced pronounced substrate-specific changes, particularly enriching several well-characterised polysaccharide-utilising species within the *Bacteroides* genus (*Bacteroides ovatus*, *Bacteroides thetaiotaomicron*, *Bacteroides uniformis*, *Bacteroides caccae*) and *Phoecaeicola vulgatus* and *Phoecaeicola dorei*. These taxa increased strongly and significantly under INU+PSY fermentation but were nearly absent in both INU and INU+MC.

**Figure 5.**
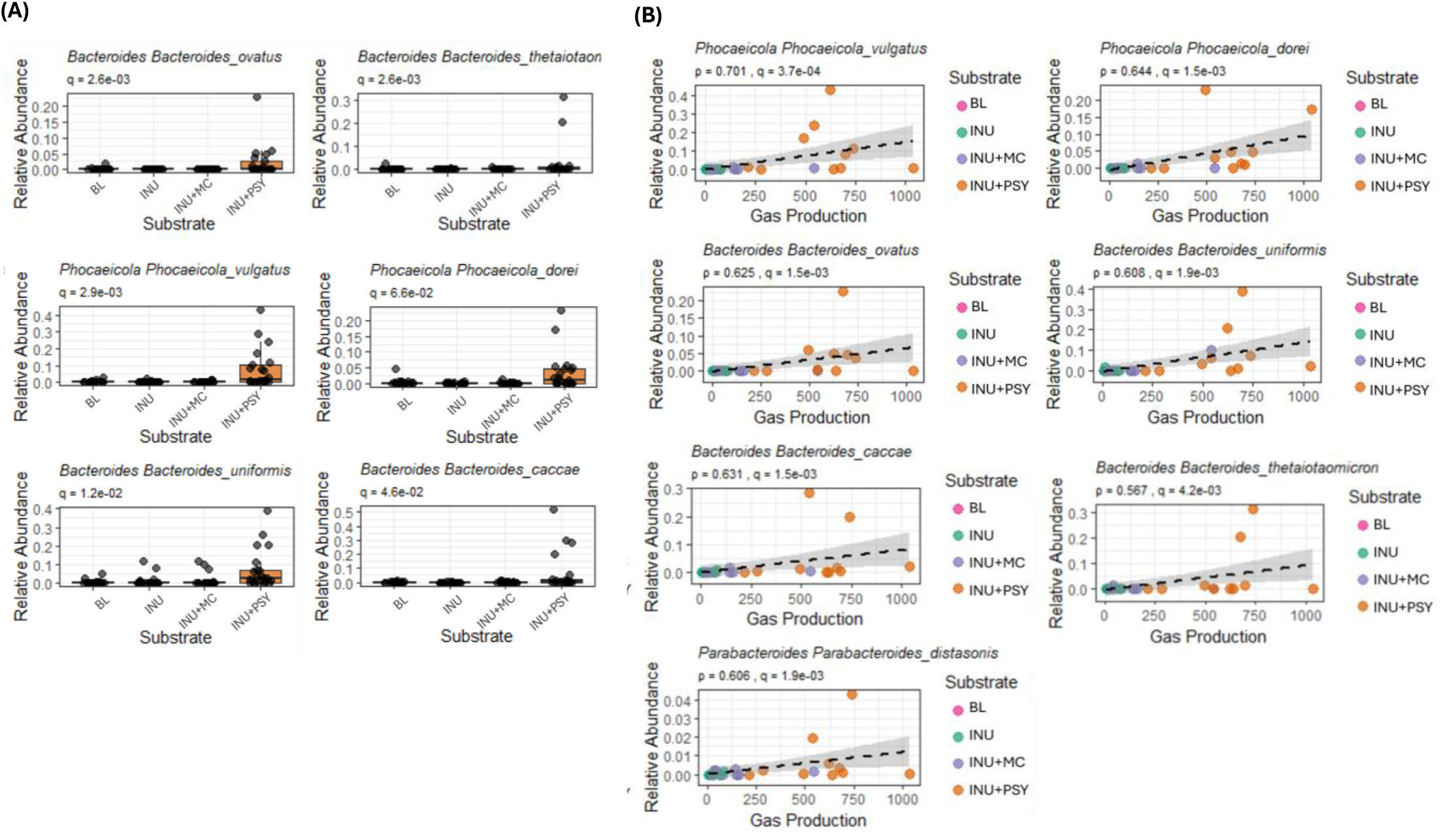
Psyllium selectively enriches key polysaccharide-utilising species. (A) Substrate-dependent differences in microbial composition at 24 hr. Relative abundance of the top 12 taxa (ranked by adjusted p-value) across substrate conditions (BL, INU, INU+MC, INU+PSY) are shown as boxplots with overlaid individual data points. Differences were assessed using the Kruskal–Wallis test with Benjamini–Hochberg false discovery rate correction (n = 48 taxa analysed; q-values shown). (B) Association between gas production and relative abundance of selected taxa. Scatter plots show relationships between total gas production and the relative abundance of the top taxa across substrates. Points represent individual samples coloured by substrate, with linear regression fits (±95% CI). Associations were assessed using Spearman’s rank correlation (ρ) with FDR-adjusted p-values (q-values).

Conversely, INU+PSY fermentation strikingly suppressed the enrichment of *Escherichia*, which was markedly increased at both 24 and 48 hr in the INU and INU+MC conditions (Fig. S4). This pronounced inhibitory effect suggests that PSY actively restrains the expansion of opportunistic taxa under these conditions, possibly reflecting its unique compositional profile, unlike INU and MC, which contain negligible protein, PSY contains approximately 2–3% protein, which, though modest in absolute terms, may represent a meaningful nitrogen source capable of shaping microbial community dynamics and competitive exclusion of opportunistic genera.

Spearman correlation between specific microbial taxa and gas production and the fermentation metabolites revealed additional mechanistic links (Fig. 6 and Table S1). *P. vulgatus* (p adj.=0.001), *P. dorei* (p adj.=0.005) and *Bacteroides* species (*B. thetaiotaomicron* (p adj.=0.01), *B. ovatus* (p adj.=0.006), *B. caccae* (p adj.=0.006), *B. uniformis* (p adj.=0.007) and were all positively associated with gas production. In contrast, *E. coli* showed a negative association with butyrate production (p adj. = 0.03), consistent with its enrichment under conditions that produce less favourable SCFA profiles. *P. vulgatus* was also positively associated with both butyrate (p. adj.=0.005) and propionate (p adj.=0.007).

**Figure 6.**
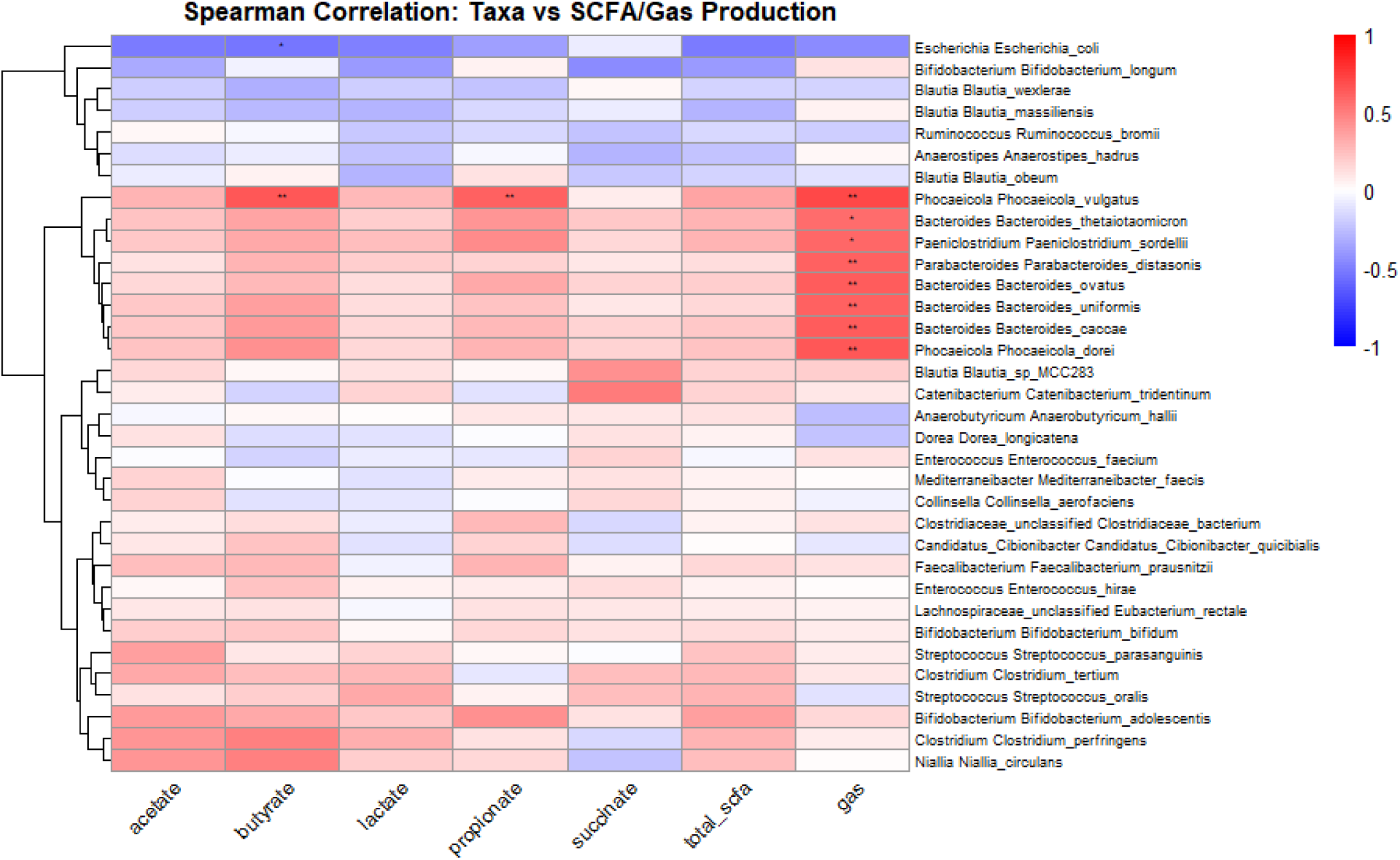
Distinct bacterial taxa are linked to fermentation outputs. Spearman correlation heatmap showing bacterial taxa co-vary with specific metabolic outputs across samples. Statistically significant correlations are depicted by * (p<0.05) and ** (p<0.01).

Taken together, these results indicate that PSY acts as a potent modulator of microbial community structure during fermentation. By selectively enriching fibre-degrading and SCFA-producing taxa while suppressing more opportunistic species, PSY may contribute to the enhanced fermentation kinetics and metabolic outputs observed above, supporting its distinct functional properties among the tested substrates.

### 4.5. Psyllium facilitates microbial degradation of fermentable fibres by providing a structural platform

We hypothesised that PSY enhances the microbial degradation of INU by providing a structural matrix that increases bacterial access to fermentable substrates. As a naturally occurring, microbially recognisable fibre, PSY may present a more favourable carbon source than the semi-synthetic MC matrix, further promoting colonisation by fibre-utilising taxa.

To investigate the spatial organisation of bacteria within fibre gels during fermentation, FISH coupled with confocal microscopy was performed on fermented PSY and MC gels (Fig. 7). Distinct differences in microbial localisation were observed between substrates, indicating that gel structure influenced bacterial distribution within the gel matrix.

**Figure 7.**
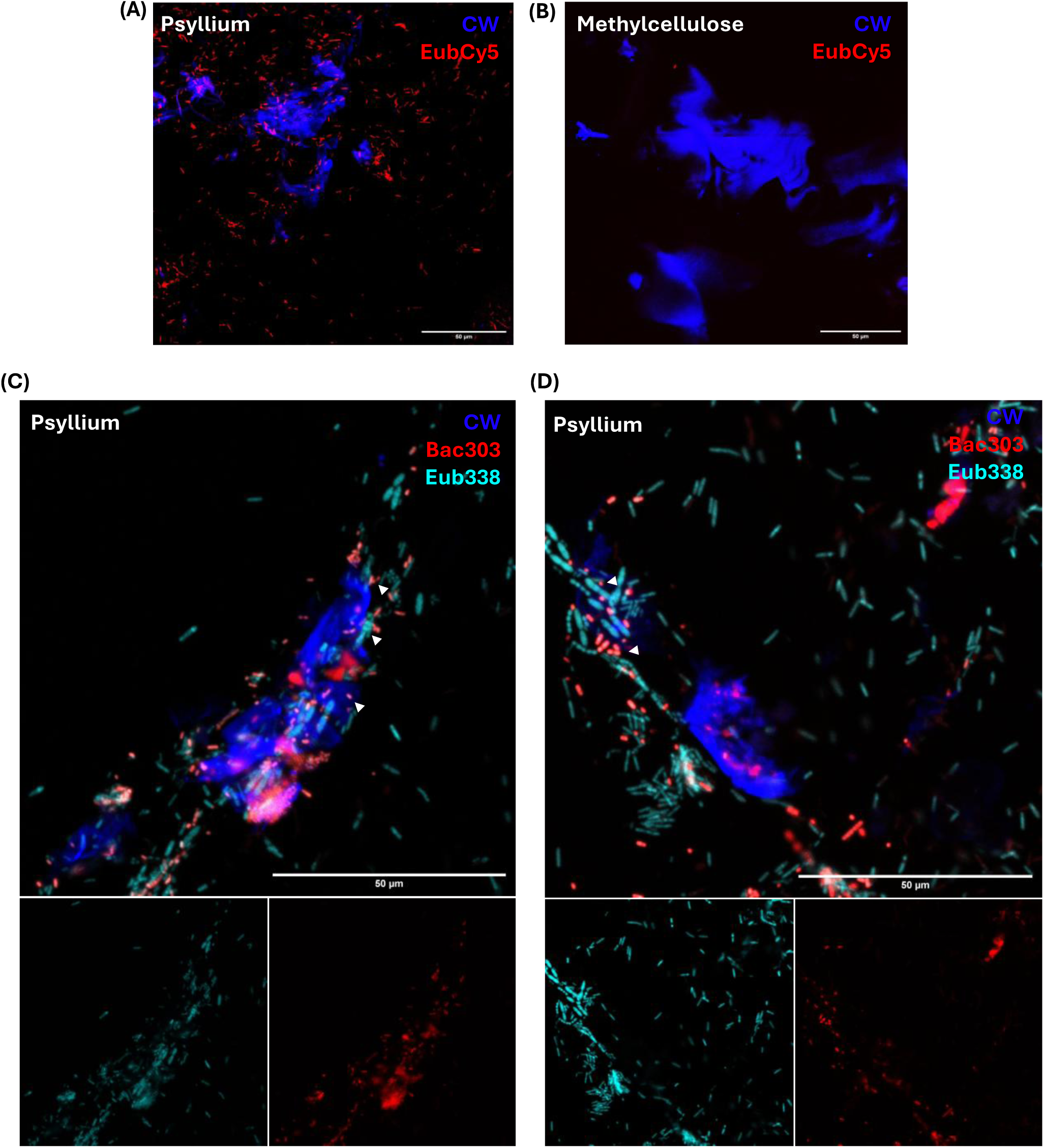
Spatial arrangement of microbiome during the fermentation of psyllium and methylcellulose. Fluorescence *in situ* hybridization (FISH) of (A) psyllium (PSY) and (B) methylcellulose (MC) gels following 24 hr *in vitro* fermentation with healthy human stool. Sections were fixed and stained with Eub338 Cy5 probe (red) and co-stained with calcofluor white (CW; blue). (C and D) Additional Bac303 TxRed probe (red) was used with Eub338 Cy5 (cyan) to determine localisation of Bacteroides species within the PSY gel-matrix. Arrowheads indicate close co-localisation of fibre and bacteria. Images were taken on confocal microscope at x20 with 2x Zoom (A and B), and x40/oil immersion x2 Zoom (C and D). Scale bars = 50 µm.

During PSY fermentation, bacteria were observed not only in the surrounding medium but also concentrated in close proximity to, and in some cases within, the PSY fibre network (Fig. 7A). This suggests active bacterial colonisation of the gel matrix, supporting enhanced access to fermentable substrates. In contrast, the MC gels showed virtually no bacterial infiltration (Fig. 7B), with cells absent from the gel matrix and surrounding medium.

Whilst the Eub338 probe identified extensive bacterial colonisation within and around the PSY matrix, the *Bacteroides*-specific probe (Bac303) revealed that taxa belonging to this genus were preferentially localised at the PSY gel interface (Fig. 7C and D). This observation is consistent with the finding that PSY leads to selective *Bacteroides* enrichment (Fig. 5), a key group of fibre-degrading bacteria.

The extensive bacterial colonisation of PSY gels may result from a combination of factors: the fibrous seed-husk structure could facilitate microbial aggregation and interaction, while specific structural motifs within the polysaccharide may be required for enzymatic recognition and fermentation. In contrast, MC lacks the chemical cues and fibrous architecture (Yu, Yakubov et al. 2017, Ren, Yakubov et al. 2020), thereby limiting bacterial penetration and metabolic activity.

### 4.6. Fermentation products of inulin and psyllium stimulate GLP-1 and 5-HT secretion in STC-1 cells

To investigate the functional impact of fermentation-derived metabolites on gut hormone secretion, murine enteroendocrine STC-1 cells were exposed to fermentation supernatants collected at 24 hrs from anaerobic batch fermentations of INU, INU+PSY, and INU+MC. Cells were subsequently immunolabelled for GLP-1 and 5-HT (Fig. 8A).

**Figure 8.**
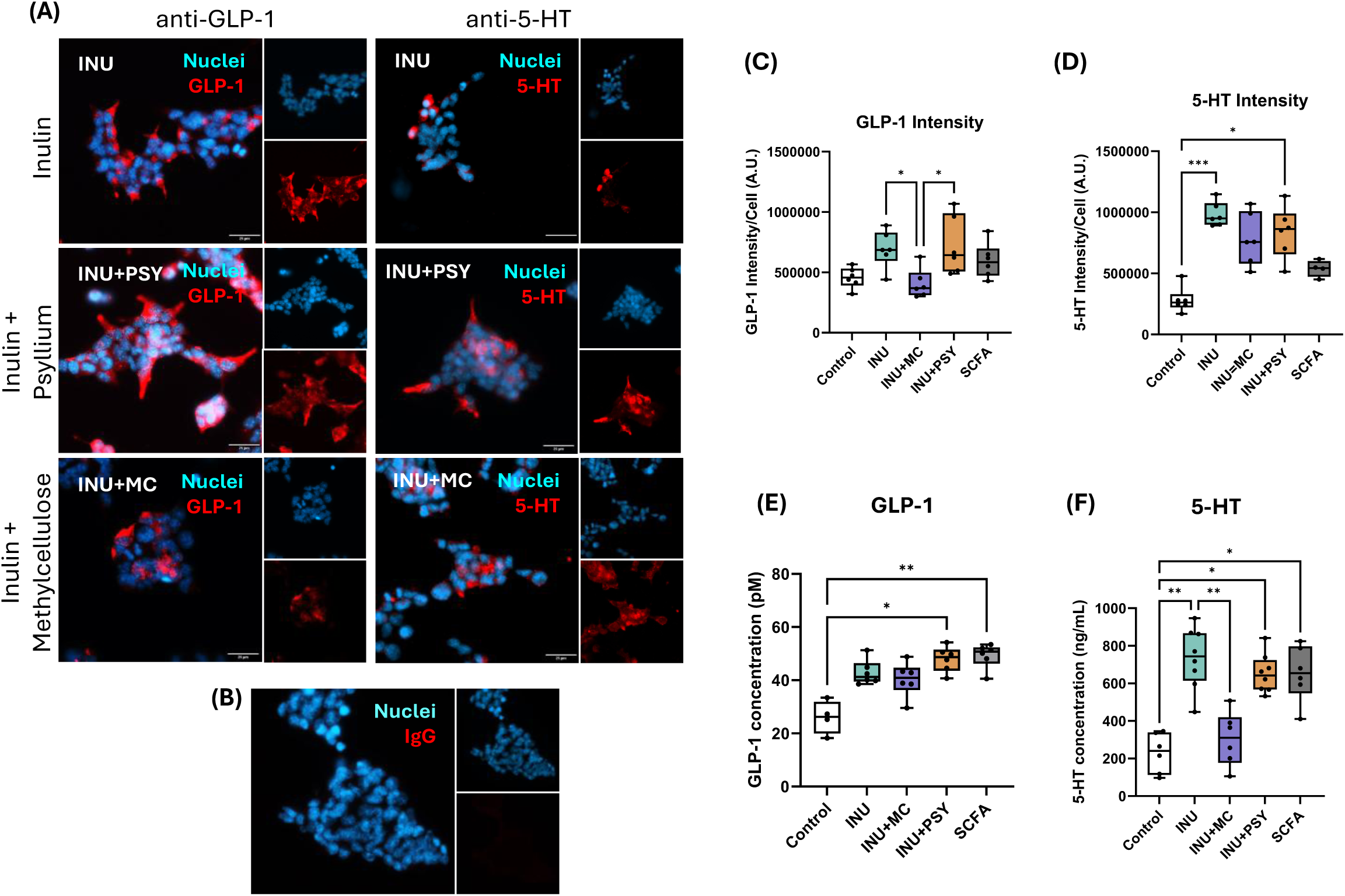
Products of inulin and psyllium fermentation stimulate GLP-1 and 5-HT secretion from STC-1 cells. STC-1 cell cultures exposed to exposed to inulin (INU; n=6), inulin+psyllium (PSY; n=6) and inulin+methylcellulose (MC; n=6) fermentation products (1:10 dilution in KREBS buffer). SCFA mix (n=6) was used as a positive control containing 10mM mixture of acetate:propionate:butyrate at a ratio of 60:20:20. Untreated cells were used as the control (n=6). Following 4 hr incubation, cells were fixed and co-stained with antibodies against GLP-1 and 5-HT. (A) Photomicrographs showing GLP-1/5-HT (red) and nuclear stain Hoechst (blue) in STC-1 cells exposed fermentation products. (B) Photomicrograph showing control IgG staining in STC-1 cells showing no immunofluorescence confirming the specificity of the antibody used. Mean staining intensities of antibodies were quantified and normalised for (C) GLP-1 and (C) 5-HT. A minimum of 10 field of view (FOV) images from each treatment group were used for calculations. (E) GLP-1 and (F) 5-HT hormone secretions from STC-1 cells exposed to fermentation. Images taken on Zeiss AxioImager Widefield Fluorescent microscope (x40/objective). Scale bar = 25 µm. Quantification data represented as mean ± SEM. One-way ANOVA followed by Tukey’s post hoc test was applied for statistical analysis (GraphPad Prism v10). Values of *p<0.05, **p<0.01 and p<0.001 were considered statistically significant.

Quantification of GLP-1 fluorescence revealed increased mean GLP-1 intensity per cell (intensity/cell; A.U.) in cells treated with fermentation products from INU+PSY fermentation (Fig. 8C). This was only statistically significant (p<0.05) compared to INU+MC fermentation (803632±94138 A.U. vs 406316±121601 A.U. for INU+PSY and INU+MC, respectively), comparable to the SCFA positive control (746116±158632 A.U.). This was reflected in a statistically significant (p<0.05) increase in GLP-1 secretion in cells exposed to INU+PSY fermentation compared to untreated cells (Fig. 8E; 47.9±1.9 pM vs 26±3.1 pM, respectively). Cells treated with INU+MC showed no significant differences in either GLP-1 intensity per cell or hormone secretion compared with untreated controls (Fig. 8C and E), indicating a lack of bioactive fermentation products capable of stimulating GLP-1 release under these conditions. Although INU and INU+PSY produced broadly comparable total SCFA profiles (Fig. 3), INU+MC fermentation did not induce GLP-1 secretion, suggesting that total SCFA abundance alone is insufficient to drive this response.

A similar trend was observed for 5-HT: intensity/cell was significantly elevated in response to INU (p<0.001) and INU+PSY fermentation (p<0.05) treatments compared to untreated controls (Fig. 8D; 982079±40412 A.U., 836388±86098 A.U. and 281575±42479 A.U. for INU, INU+PSY and BL, respectively). This was reflected in significant increases in 5-HT secretion, in response to both INU and INU+PSY fermentation products compared to untreated cells (Fig. 8F).

INU+MC-treated cells showed no statistically significant difference in either intensity/cell or hormone secretion from untreated controls for both GLP-1 and 5-HT (Fig.8C-F), suggesting that total SCFA levels alone do not determine bioactivity of fermentation supernatants.

These findings indicate that fermentation products derived from INU and PSY, particularly in combination, possess bioactivity capable of stimulating gut hormone secretion. In contrast, INU+MC-derived fermentation products did not stimulate hormone release despite producing SCFA levels comparable to those produced by INU alone, suggesting a lack of stimulatory bioactivity rather than active inhibition. This supports the hypothesis that gut hormone secretion is influenced by specific SCFA profiles or additional microbial metabolites, rather than total SCFA abundance. Further mechanistic studies are needed to confirm the potential inhibitory effect of MC on EEC secretion.

## 5. Discussion

Our findings appear paradoxical at first glance. PSY accelerated INU fermentation and increased gas production in our controlled *in vitro* system (Fig. 2), whereas previous clinical studies demonstrated that PSY co-administration delayed gas production and reduced IBS symptoms in human participants (Gunn, Abbas et al. 2022, Alhasani, Modasia et al. 2024, Reid, Alhasani et al. 2026). This discrepancy may be an apparent manifestation of the spatial and temporal dynamics of colonic fermentation that are present *in vivo* but absent in static batch fermentation models.

In the gut, PSY’s high water-holding capacity and viscoelasticity slow gastric emptying and intestinal transit (Major, Murray et al. 2018), delaying arrival in the colon. It also has an inhibitory effect within the colon, delaying colonic fermentation, shifting the site of fermentation distally along the colon. (Alhasani, Modasia et al. 2024). This redistribution, rather than a reduction in overall fermentation, is likely central to its clinical benefits (So, Yao et al. 2023, Bertin, Zanconato et al. 2024). Furthermore, the reported self-healing characteristics of PSY hydrogels would explain their ability to maintain gel structure through gastrointestinal transit (Yu, Stokes et al. 2021). Displacing the fermentation distally into the transverse and descending colon could reduce proximal colon distension thereby alleviating symptoms while maintaining or enhancing SCFA production.

Our *in vitro* system lacks transit dynamics and instead reflects events once the PSY–INU matrix reaches the colon. The sustained increase in gas and SCFA production (Fig. 2 and 3), therefore, demonstrates that PSY does not restrict microbial access to INU but instead facilitates fermentation (Fig. 7). This interpretation aligns with clinical data showing that PSY supplementation maintains or increases faecal SCFA concentrations while improving IBS symptoms (Jalanka, Major et al. 2019, Gunn, Abbas et al. 2022, Garg, Garg et al. 2024). The consistent elevation of lactate (Fig. 3D) across replicate experiments suggests enhanced early-stage fermentation activity during co-fermentation with PSY, however, substantial inter-experimental variability likely limited the statistical resolution between treatments. These precursor metabolites are commonly produced during the initial stages of fermentation and can serve as substrates for cross-feeding bacteria to generate downstream SCFAs (Macfarlane and Macfarlane 2003). The accumulation of lactate in fermentation, and increases in acetate, propionate, and butyrate, suggest enhanced microbial metabolic flux and more efficient fermentation dynamics when PSY is co-fermented with INU. In contrast, INU+MC fermentation did not significantly enhance or diminish SCFA production relative to INU alone (Fig. 3). This indicates that PSY selectively enhances microbial metabolism of INU, resulting in greater production of beneficial fermentation metabolites, whereas MC does not.

The delayed onset of isolated PSY degradation in our system (Fig. S1), beginning at ∼20 hrs, further supports this interpretation. PSY remains relatively resistant to early degradation, consistent with its known slow fermentability (Marteau, Flourié et al. 1994, McRorie 2013, Harris, Pereira et al. 2023). However, rather than acting as an inert barrier, PSY creates a microenvironment that enhances bacterial access to co-administered fermentable substrates. Imaging revealed marked differences in microbial spatial organisation between PSY and MC (Fig. 7). Gels prepared from PSY supported extensive bacterial colonisation, with microbes not only adhering to fibre surfaces but also penetrating the gel matrix and forming dense clusters. In contrast, MC gels remained largely devoid of bacterial infiltration despite similar bulk rheological properties. This may be due to the methoxylation of hydroxyl groups present on MC monomer units, creating more inert surface in stark contrast to the fibre surface of PSY. This indicates that gelation alone is insufficient to promote microbial colonisation and that fibre chemistry plays a decisive role.

These differences likely reflect fundamental variation in bacterial recognition and utilisation of the fibre substrates. Unlike cellulose or MC, which are relatively structurally uniform polymers, PSY comprises a heterogeneous mixture of complex, highly substituted polysaccharides, predominantly arabinoxylans. These arabinoxylans consist of a β-1,4-linked xylose backbone variably substituted with arabinose side chains, resulting in substantial structural diversity that can influence microbial accessibility and fermentation dynamics (Izydorczyk and Biliaderis 1995, Yu, Yakubov et al. 2017, Ren, Yakubov et al. 2020). This structural motif is readily recognised by polysaccharide utilisation loci (PULs) present in many gut bacteria, particularly members of the Bacteroidetes phylum (Martens, Lowe et al. 2011, El Kaoutari, Armougom et al. 2013, Drula, Garron et al. 2022). PULs encode the enzymatic machinery for polysaccharide binding, depolymerisation, and degradation, enabling bacteria to identify and colonise compatible polysaccharides In contrast, MC, a common cellulose semi-synthetic derivative, consists of glucose units connected by β-1,4-glycosidic bonds with methyl group substitutions that render it resistant to microbial degradation (Machle, Heyroth et al. 1944, Ferguson and Jones 2000). Lacking the structural motifs, MC fails to recruit colonising bacteria despite forming a gelling matrix.

Consistent with this mechanism, INU+PSY fermentation selectively enriches polysaccharide-degrading taxa including *Bacteroides* and *Phoecaecoli* species (Fig. 5). *B. thetaiotaomicron* harbours over 80 distinct PULs, (Porter, Luis et al. 2018) while *B. ovatus*, *B. uniformis*, and *B. caccae* similarly possess extensive carbohydrate-degrading capabilities (Martens, Lowe et al. 2011, Rakoff-Nahoum, Coyne et al. 2014, Singh, Rajarammohan et al. 2020). *B. ovatus* is particularly notable for its ability to utilise INU directly or via glycoside hydrolases secretion (Rakoff-Nahoum, Foster et al. 2016). The selective expansion of these taxa suggests that PSY acts as both a colonisable scaffold and a co-substrate. This spatial organisation likely enhances cross-feeding interactions and metabolic efficiency (Macfarlane and Macfarlane 2003, Cronin, Joyce et al. 2021). In contrast, INU+MC and INU were similar, indicating that, in these *in vitro* experiments, the non-fermentable nature of MC limited microbial growth, and the physical properties of the gels were less influential than PSY’s ability to act as both a scaffold and a co-substrate.

This interpretation is further supported by strong positive correlations between *P. vulgatus* (formerly *Bacteroides vulgatus*) abundance and both gas production and SCFA production (Fig. 5 and S1) . *P. vulgatus* is a key cross-feeder, consuming fermentation intermediates such as lactate and acetate produced by primary fermenters to generate propionate and butyrate (Kattel, Morell et al. 2023, Solvang, Farquharson et al. 2025). The PSY gel matrix may promote such syntrophic interactions by maintaining bacteria in close spatial proximity, facilitating efficient metabolic transfer.

INU+PSY fermentation also showered higher Shannon and Simpson diversity compared with other conditions (Fig. 4A and B). Fermentable substrates typically reduce diversity during early fermentation as fast-growing specialists dominate (Cantu-Jungles and Hamaker 2023). The maintenance of diversity with PSY likely reflects the structural heterogeneity of the gel matrix, which provides multiple ecological niches that can support a broader range of taxa, rather than differences in fermentation kinetic (Harris, Pereira et al. 2023).

The functional relevance of these microbial changes was underscored by the enhanced secretion of GLP-1 and serotonin (5-HT) by STC-1 cells exposed to INU+PSY fermentation supernatants (Fig. 8). GLP-1 plays keys roles in regulating intestinal secretion, motility, and glucose homeostasis (Amato, Cinci et al. 2010, Baggio and Drucker 2014) while 5-HT regulates and modulates gut motility, secretion, and visceral sensation (Bornstein 2012, Mawe and Hoffman 2013). The enhanced GLP-1 secretion was comparable to that induced by SCFA-positive controls, strongly suggesting SCFA-mediated mechanisms. Propionate and butyrate stimulate GLP-1 release via free fatty acid receptors free fatty acids receptor 2 (FFAR2/GPR43) and FFAR3 (GPR41) expressed on EECs (Tolhurst, Heffron et al. 2012). The elevated propionate and butyrate concentrations following INU+PSY fermentation provide a plausible mechanism for the observed hormone responses. While SCFAs are likely primary drivers, they do not fully explain the higher hormone secretion seen with INU compared with INU+MC, despite similar SCFA concentrations (Grant, De Franco et al. 2025). One possibility is that the physical or chemical properties of MC, such as the methoxyl groups creating hydrophobic regions, may limit SCFA release from the gel matrix, particularly for more hydrophobic SCFAs like propionate and butyrate. Other fermentation-derived metabolites may also contribute, highlighting the need for future metabolomic investigation (Verbeke, Boobis et al. 2015).

These findings have important clinical implications. Altered enteroendocrine signalling is implicated in IBS pathophysiology (El-Salhy and Gilja 2017) and appropriate GLP-1 and 5-HT signalling may reduce visceral sensitivity and discomfort (O’Brien and O’Malley 2020, Mostafa and Alrasheed 2025). The ability of INU+PSY-derived fermentation products to stimulate these hormones suggests that PSY’s benefits arise from multiple complementary mechanisms including spatial redistribution of fermentation, enhancement of SCFA production and modulation of gut-microbe-host signalling. MC, despite similar physical properties including delayed release of INU *in vitro* (Reid et al 2026), failed to engage this microbial–metabolic–hormonal axis.

In conclusion, this study challenges the assumption that gel-forming fibres slow fermentation through physical entrapment of fermentable substrates. Instead, we show that gel-forming fibre PSY actively enhances INU fermentation *in vitro* through mechanisms dependent on its fermentability and microbial compatibility rather than on gelation alone. Notably, PSY co-fermentation was accompanied by the formation of a stable foam with numerous small bubbles, an observation absent in MC fermentations, which may have important *in vivo* implications. Specifically, foam-mediated gas entrapment within the gut could delay the appearance of fermentation gases in breath tests, potentially leading to an underestimation of PSY fermentability in clinical hydrogen breath test studies despite active and robust fermentation occurring. By comparing PSY with MC, a cellulose-derivative with similar physiochemical properties but lacks fermentability, we provide direct evidence that the biological activity of gel-forming fibres, and not just their physical structure, determines their impact on colonic fermentation dynamics and host-microbe signalling. We show that PSY acts as a colonisable scaffold, concentrating bacteria near fermentable substrates, selectively enriching SCFA-producing taxa, and generating metabolites capable of stimulating gut hormone secretion. In contrast, MC seemed to exclude bacteria from the fibre gel surface and did not stimulate gut hormone secretions, suggesting that the chemically modified structure of MC may impact biological compatibility, rationalising the shortcomings of MC in terms of SCFA production. These findings highlight a mechanistic basis for the beneficial effects of PSY on gut microbial ecology and signalling, with potential relevance for dietary strategies aimed at supporting gut health and alleviating symptoms in functional gastrointestinal disorders such as IBS.

## Acknowledgement

The authors gratefully acknowledge the support of the QIB Colon Model Facility; the facility was funded by the BBSRC Core Capability Grant BB/CCG2260/1 and BBSRC FMH ISP Grant BB/X011054/1.

The authors also thank the participants in the QIB Colon Model Study (IRAS 171172) for donating stool samples.

The authors thank Kathryn Gotts of Quadram Institute Biosciences Advanced Microscopy Facility for their support & assistance in this work and the Sample Preparation Facility which is funded by the BBSRC Investment Gateway Panel IGP24-11.

## Data availability statement

The data supporting the findings of this study are available from the corresponding author upon reasonable request.

