## Supplementary Figure 1 for "Gel-forming fibres differentially modulate inulin fermentation: A comparison of psyllium and methylcellulose in *in vitro* colonic models"

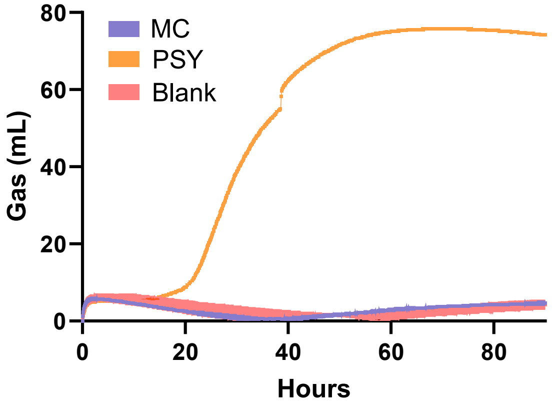


**Supplementary Figure 1. *In vitro* psyllium is fermentable whilst methylcellulose is not.** Gas production kinetics assessed from fresh healthy donor stools (n=1-2) using the ANKOM RF system reveals that psyllium (PSY) is fermentable after ~20 hr during *in vitro* anaerobic fermentation, compared to methylcellulose (MC) that remains non-fermentable. No substrate (BL) was used as a control.
