## Supplementary Figure 2 for "Gel-forming fibres differentially modulate inulin fermentation: A comparison of psyllium and methylcellulose in *in vitro* colonic models"

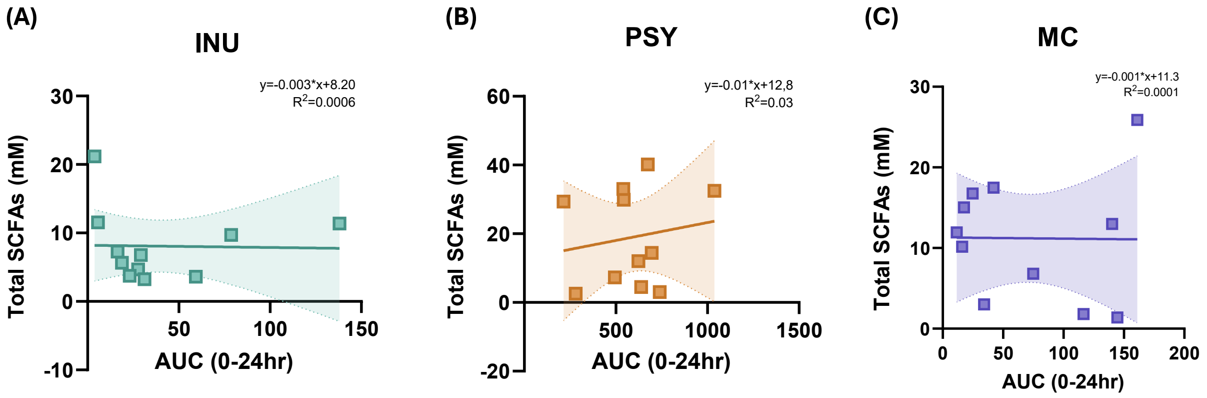


**Supplementary Figure 2. Correlation between *in vitro* SCFA production and gas at 24 hr fermentation**. Graphs showing relationship between total SCFA production (mM) and *in vitro* gas area under the curve (AUC) at 24 hr fermentation of (A) inulin (INU), (B) inulin+psyllium (PSY) and (C) inulin+methylcellulose (MC). Simple Linear regression was calculated on GraphPad (GraphPad Prism v10).
