## Supplementary Figure 3 for "Gel-forming fibres differentially modulate inulin fermentation: A comparison of psyllium and methylcellulose in *in vitro* colonic models"

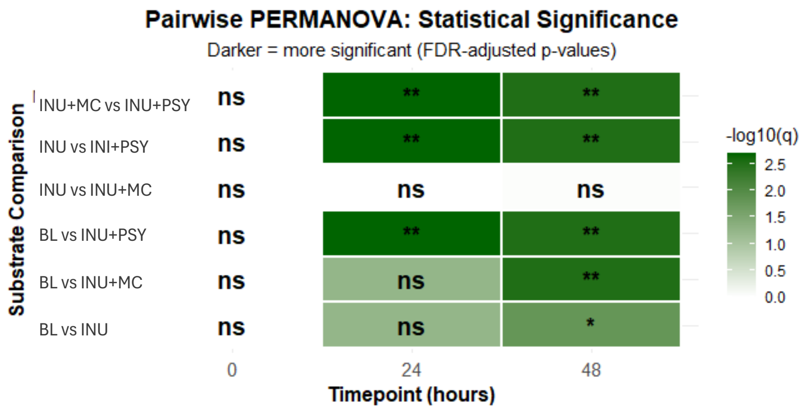


**Supplementary Figure 3. Microbial community composition differs between substrates over time.** Summary heatmap of pairwise PERMANOVA results over 0, 24 and 48 hr timepoints for each fibre treatment; *p<0.05, **p<0.01.
