## Supplementary Figure 4 for "Gel-forming fibres differentially modulate inulin fermentation: A comparison of psyllium and methylcellulose in *in vitro* colonic models"

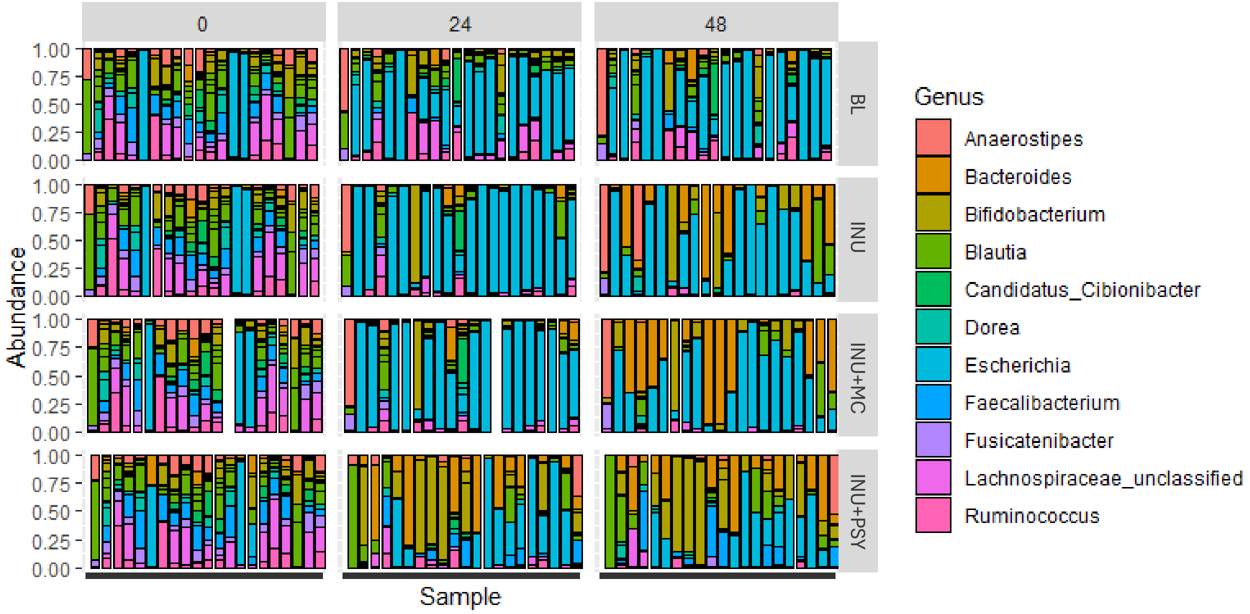


**Supplementary Figure 4. Psyllium differentially modulates microbial composition.** Abundance plots showing genus level abundances across different fermentation timepoints (0, 24 and 48 hr) for each substrate (inulin alone (INU) and co-fermented with psyllium (INU+PSY) and methylcellulose (INU+MC). No substrate used as control (BL).
