## Supplementary Table 1 for "Gel-forming fibres differentially modulate inulin fermentation: A comparison of psyllium and methylcellulose in *in vitro* colonic models"

| Species | Variable | Correlation | P_value | P_adjusted |
|---|---|---|---|---|
| Phocaeicola_vulgatus | gas | 0.700913 | 7.91E-06 | 0.001883 |
| Phocaeicola_vulgatus | butyrate | 0.652609 | 5.17E-05 | 0.005534 |
| Phocaeicola_dorei | gas | 0.644007 | 6.98E-05 | 0.005534 |
| Bacteroides_ovatus | gas | 0.625229 | 0.00013 | 0.0062 |
| Bacteroides_caccae | gas | 0.630585 | 0.000109 | 0.0062 |
| Phocaeicola_vulgatus | propionate | 0.613658 | 0.000188 | 0.0071 |
| Bacteroides_uniformis | gas | 0.608207 | 0.000222 | 0.0071 |
| Parabacteroides_distasonis | gas | 0.605785 | 0.000239 | 0.0071 |
| Paeniclostridium_sordellii | gas | 0.584635 | 0.000442 | 0.011679 |
| Bacteroides_thetaiotaomicron | gas | 0.567105 | 0.000713 | 0.016972 |
| Escherichia_coli | butyrate | -0.53152 | 0.001745 | 0.037753 |

**Supplementary Table 1. Significantly associated bacterial taxa from Spearman correlation.**
